# Refinement of Multiconformer Ensemble Models from Multi-temperature X-ray Diffraction Data

**DOI:** 10.1101/2023.05.05.539620

**Authors:** Siyuan Du, Stephanie A. Wankowicz, Filip Yabukarski, Tzanko Doukov, Daniel Herschlag, James S. Fraser

**Affiliations:** Department of Biochemistry, Stanford University, Stanford, California 94305, United States; Department of Chemistry, Stanford University, Stanford, California 94305, United States; Department of Bioengineering and Therapeutic Sciences, University of California, San Francisco, San Francisco, California 94143, United States; Bristol-Myers Squibb, San Diego, California 92121, United States; Stanford Synchrotron Radiation Lightsource, SLAC National Accelerator Laboratory, Menlo Park, California 94025, United States; Department of Chemical Engineering, Stanford University, Stanford, California 94305, United States; Stanford ChEM-H, Stanford University, Stanford, California 94305, United States; Quantitative Biosciences Institute, University of California, San Francisco, California 94143, United States

**Keywords:** multi-temperature X-ray crystallography, room temperature X-ray, conformational ensembles, multiconformer model, structural model refinement

## Abstract

Conformational ensembles underlie all protein functions. Thus, acquiring atomic-level ensemble models that accurately represent conformational heterogeneity is vital to deepen our understanding of how proteins work. Modeling ensemble information from X-ray diffraction data has been challenging, as traditional cryo-crystallography restricts conformational variability while minimizing radiation damage. Recent advances have enabled the collection of high quality diffraction data at ambient temperatures, revealing innate conformational heterogeneity and temperature-driven changes. Here, we used diffraction datasets for Proteinase K collected at temperatures ranging from 313 to 363K to provide a tutorial for the refinement of multiconformer ensemble models. Integrating automated sampling and refinement tools with manual adjustments, we obtained multiconformer models that describe alternative backbone and sidechain conformations, their relative occupancies, and interconnections between conformers. Our models revealed extensive and diverse conformational changes across temperature, including increased bound peptide ligand occupancies, different Ca^2+^ binding site configurations and altered rotameric distributions. These insights emphasize the value and need for multiconformer model refinement to extract ensemble information from diffraction data and to understand ensemble-function relationships.

## 1 Introduction

All molecular processes are defined by energy landscapes, which are in turn manifested by an ensemble of interconverting conformational states (Austin et al., 1975; Benkovic et al., 2008; Benkovic and Hammes-Schiffer, 2003; Frauenfelder et al., 1991, 1988; Hammes et al., 2011). For example, ligand binding affinity is defined by the relative population of the bound to the unbound state(s), and enzymatic rates by the possibility of crossing to the transition state from the ground state. Therefore, understanding protein functions requires obtaining and comparing conformational ensembles in different bound states under physiologically-relevant conditions. Because conformational ensembles reveal probabilities of states and therefore their underlying energetics, they provide the possibility to relate structural features to thermodynamic quantities for molecular processes – a goal unattainable using single conformer structural models and an essential step towards a quantitative and predictive understanding of protein functions.

The need for conformational ensembles to decipher protein functions has long been recognized, yet experimental approaches to obtain ensemble information are limited by their resolution or by technological challenges. For example, nuclear magnetic resonance (NMR) methods allow us to determine the degree of motion of protein groups and the rate of interconversions between sub-states but do not reveal atomic-level details of these sub-states (Ishima and Torchia, 2000; Kempf and Loria, 2003; Kleckner and Foster, 2011; Kovermann et al., 2016; Mittermaier and Kay, 2006). Similarly, Förster resonance energy transfer (FRET) experiments are used to study protein conformational dynamics, but only reveal large conformational changes reported by the changes in two groups (the donor and the acceptor) (Mazal and Haran, 2019; Okamoto and Sako, 2017; Schuler and Eaton, 2008). In contrast, X-ray crystallography provides atomic-level information about protein three-dimensional structures.

The ability to model individual atom positions from diffraction data has allowed us to relate the shape of a protein to its function (Indiani and O’Donnell, 2006; Kato et al., 2018), identify specific residues involved in biological processes and propose models for how they function (Robertus et al., 1972; Tsukada and Blow, 1985). In principle, X-ray diffraction data represent an ensemble average from multiple conformational states (DePristo et al., 2004; Rejto and Freer, 1996; Smith et al., 1986), but obtaining and modeling ensembles from X-ray data have been challenging for two practical reasons. First, the majority of PDB-deposited structures are obtained under cryogenic conditions (∼100 K) (Garman, 2003). While useful in reducing radiation damage, cryo-cooling alters the conformational landscape of a protein because the low temperature strongly favors low enthalpy states and quenches many degrees of freedom (Frauenfelder et al., 1979; Weik and Colletier, 2010). As shown in multiple studies, protein dynamics typically undergo a significant change (termed “glass transition”) at ∼180 to 200 K, suggesting that structural features from models obtained under this temperature range may reflect cryo-artifacts instead of physiologically-relevant protein features (Fraser et al., 2011; Halle, 2004; Keedy et al., 2014; Rasmussen et al., 1992; Tilton et al., 1992). Indeed, crystallographic data obtained at ambient temperatures reveal conformational states that are hidden or different from cryo structures (Fraser et al., 2011, 2009; Keedy et al., 2015b; Yabukarski et al., 2022).

Second, most of the structures deposited in the PDB are modeled as single conformers, which in many cases do not explain the full density data (Smith et al., 1986). Single conformer models typically use isotropic or anisotropic B-factors to represent variability of atomic positions, but these parameters can only account for harmonic deviations from the average positions, with the assumption that atoms fluctuate within a single local minimum. Nevertheless, the protein conformational landscape is rugged, where a large fraction of residues may be able to occupy multiple local minimums of similar energies, resulting in anharmonic electron density distributions (Kuriyan et al., 1986). More recently, modeling techniques have emerged to model anharmonic displacements from the underlying diffraction data to reveal the alternative conformations that the protein can adopt (Burnley et al., 2012; Burnley and Gros, 2013; Forneris et al., 2014; Fraser et al., 2011; Ginn, 2021; Keedy et al., 2015a; Riley et al., 2021; van Zundert et al., 2018). However, unlike methods to obtain single conformer models which have become standardized and widely-applied, methods to efficiently search for and model alternative conformations require specialized software and techniques that are only used by a relatively small community.

To address these challenges in obtaining ensemble models via X-ray crystallography, we recently described an improved data collection pipeline to minimize radiation damage at ambient temperatures (up to 363K) that can be broadly implemented for different proteins and at other beamlines (Doukov et al., 2020). Here, we focus on the refinement of X-ray diffraction data obtained at ambient temperatures to generate multiconformer ensemble models of high quality and interpretability. Using diffraction datasets of Proteinase K collected at a series of temperatures (313 to 363 K) above the glass-transition range, we provide a practical roadmap to guide multiconformer model refinement and discuss refinement choices and their advantages and limitations. In addition, in these datasets across temperature, we observed changes in the binding positions of a Ca^2+^ ion that is required for catalysis, and we describe the modeling and refinement of alternative Ca^2+^ binding configurations and coupled motions of Ca^2+^-coordinating residues.

Finally, we show the profound impact of temperature on the Proteinase K conformational ensemble revealed by our models, including changes in conformational heterogeneity (such as altered rotamer distributions) and compositional heterogeneity (such as increased peptide-bound states at higher temperatures), emphasizing the need for ambient- and multi-temperature X-ray crystallography to probe protein conformational landscapes and reveal hidden conformational features.

## 2 Collection of multi-temperature X-ray diffraction data

### 2.1 Obtaining crystals for X-ray diffraction at and above room temperature

*Tritirachium album* proteinase K (Sigma, catalog # P2308) was dissolved at pH 7.5 to 30 mg/mL in a 50 mM TRIS (Sigma, T1699) buffer . The protein was crystallized using a hanging drop setup on a 24 well VDX plate with sealant (Hampton Research, HR3-171) and 22 mm thick siliconized circle cover slides (Hampton Research, HR3-247) by mixing 2 µL of protein solution with 2 µL 1.2 M ammonium sulfate, AS (Sigma, A4915) on the coverslip, which was placed over 1 mL 1.2 M AS in the VDX plate well. Prior to data collection, the aqueous layer around the crystals was exchanged to an inert Paratone-N oil (Hampton Research; # HR2-643).

Paratone-N oil layer significantly reduces evaporation (Hope, 1990; Pflugrath, 2015; Weik et al., 2005). Oil-exchanged crystals were mounted on Dual-Thickness MicroLoops LD™ (Mitegen, SKU:M2-L18SP-200) and MicroGrippers™ loops (Mitegen, SKU:M7-L18SP-300). Excessive oil was removed, and pins were manually mounted on the BL14-1 goniometer at Stanford Synchrotron Radiation Lightsource (SSRL) for data collection (Doukov et al., 2020). Additional information on the crystallization protocol can be found at https://www.moleculardimensions.com/products/ready-to-grow-crystallization-kit.

### 2.2 Achieving high-temperature capabilities and temperature control

An Oxford Cryosystems Cryostream 800 model N2 cooler/heater (https://www.oxcryo.com/single-crystal-diffraction/cryostream-800) with a temperature range of 80-400 K was installed to collect high temperature data at the SSRL beamline 14-1. Because the physical properties of protein crystals deteriorate over time when exposed to high temperatures, we adapted the standard nozzle-closing crystal annealer operation to control the crystal exposure to the heated N_2_ stream and minimize time at high temperature as follows. After the N_2_ gas is heated to the desired (high) temperature, the annealer paddle is placed in the “closed” position to prevent the gas flow from reaching the sample and heating it during the experimental setup [i.e., crystal mounting and centering, closing the experimental hutch, entering the experimental parameters into the *Blu-Ice* control software (McPhillips et al., 2002)]. Control kinetic measurements showed that a J thermocouple placed from room temperature (∼293 K) to a 363 K N2 stream (the highest temperature used in this work) was within 5% of the desired temperature in ≤ 10 seconds (not shown) and we used this equilibration time prior to data collection (see below). For data collection, the annealer paddle is moved to the “open” position via the beamline control software *Blu-Ice* and data collection is initiated after a ≤10 seconds temperature equilibration delay (Doukov et al., 2020).

### 2.3 Diffraction data collection

Proteinase K crystals with dimensions 0.3-0.4 mm on each side were used for data collection. Larger crystals are required for the collection of X-ray diffraction data at and above room temperature to approach cryo resolutions, because higher temperature can lead to more radiation damage (Garman and Owen, 2006; Garman and Weik, 2017; Nave and Garman, 2005; Roedig et al., 2016; Southworth-Davies et al., 2007; Warkentin et al., 2011; Warkentin and Thorne, 2010). To maximize diffraction intensity while minimizing the number of absorbed photons per unit cell, the beam and crystal size are matched as closely as possible. We routinely used the highest beam size of 250 µm (horizontal) by 80 µm (vertical). At least 100 degrees of rotation data were collected as quickly as possible for each crystal to avoid dehydration and any macroscale defects in the crystal that can happen alongside microscopic radiation damage.

Usually each degree frame was collected for 0.04 – 0.2 seconds with the detector distance and energy adjusted to achieve highest resolution and high quality dataset (see Table 1, Table S1 from Doukov et al., 2020).

**Table 1.**
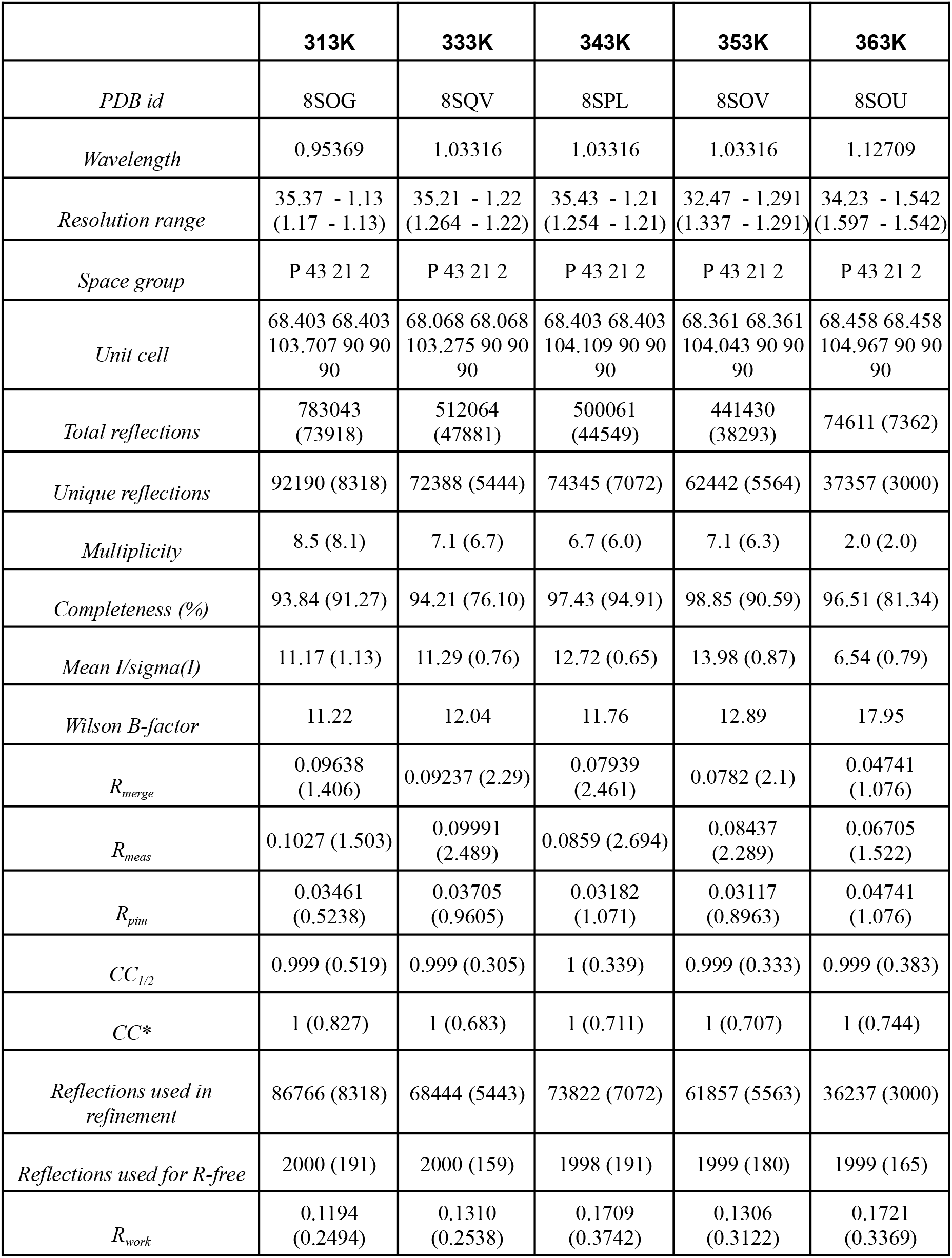
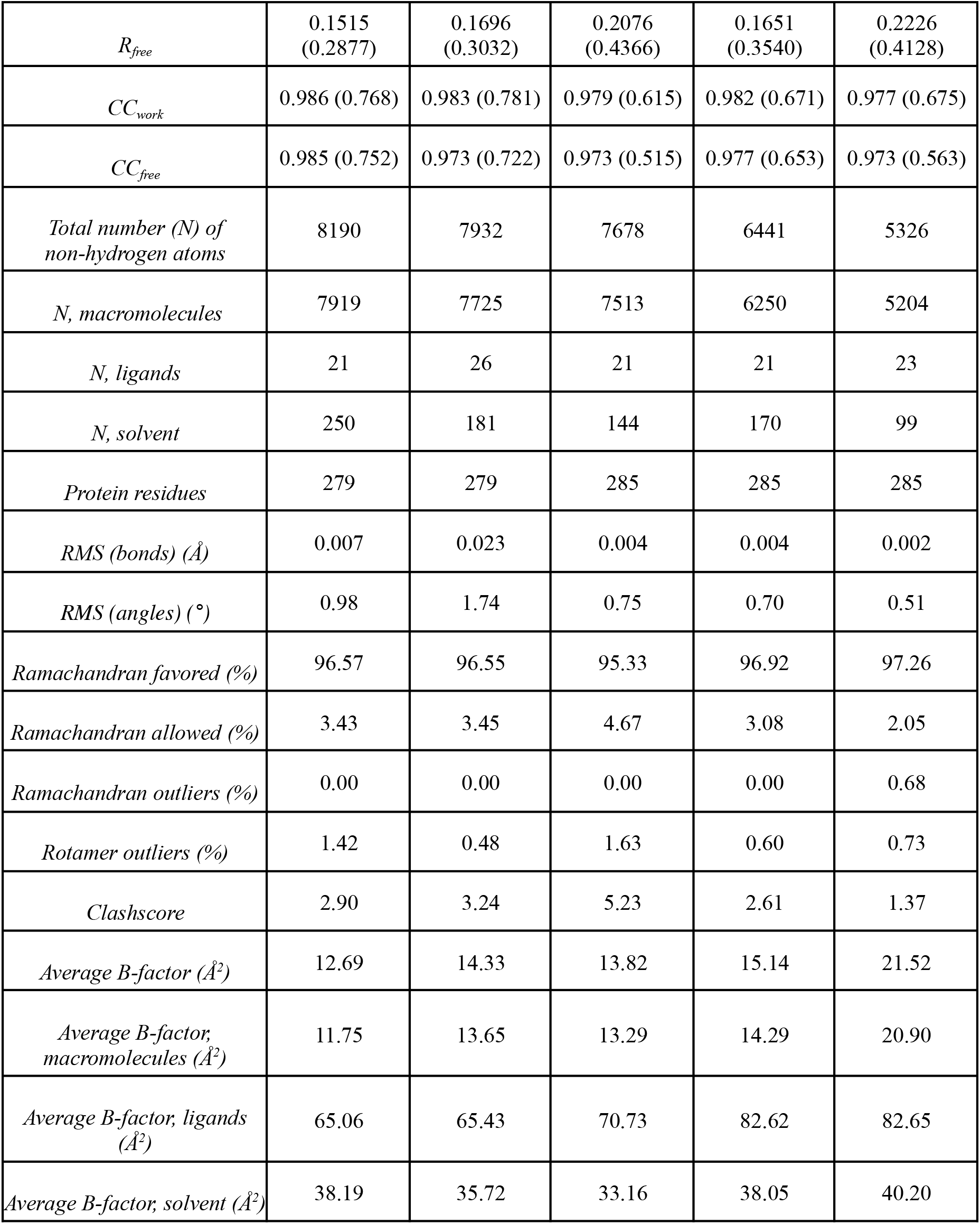
Data collection and refinement statistics. Statistics for the highest-resolution shell are shown in parentheses.

|  | **313K** | **333K** | **343K** | **353K** | **363K** |
| --- | --- | --- | --- | --- | --- |
| *PDB id* | 8SOG | 8SQV | 8SPL | 8SOV | 8SOU |
| *Wavelength* | 0.95369 | 1.03316 | 1.03316 | 1.03316 | 1.12709 |
| *Resolution range* | 35.37 - 1.13<br>(1.17 - 1.13) | 35.21 - 1.22<br>(1.264 - 1.22) | 35.43 - 1.21<br>(1.254 - 1.21) | 32.47 - 1.291<br>(1.337 - 1.291) | 34.23 - 1.542<br>(1.597 - 1.542) |
| *Space group* | P 43 21 2 | P 43 21 2 | P 43 21 2 | P 43 21 2 | P 43 21 2 |
| *Unit cell* | 68.403 68.403<br>103.707 90 90<br>90 | 68.068 68.068<br>103.275 90 90<br>90 | 68.403 68.403<br>104.109 90 90<br>90 | 68.361 68.361<br>104.043 90 90<br>90 | 68.458 68.458<br>104.967 90 90<br>90 |
| *Total reflections* | 783043<br>(73918) | 512064<br>(47881) | 500061<br>(44549) | 441430<br>(38293) | 74611 (7362) |
| *Unique reflections* | 92190 (8318) | 72388 (5444) | 74345 (7072) | 62442 (5564) | 37357 (3000) |
| *Multiplicity* | 8.5 (8.1) | 7.1 (6.7) | 6.7 (6.0) | 7.1 (6.3) | 2.0 (2.0) |
| *Completeness (%)* | 93.84 (91.27) | 94.21 (76.10) | 97.43 (94.91) | 98.85 (90.59) | 96.51 (81.34) |
| *Mean I/sigma(I)* | 11.17 (1.13) | 11.29 (0.76) | 12.72 (0.65) | 13.98 (0.87) | 6.54 (0.79) |
| *Wilson B-factor* | 11.22 | 12.04 | 11.76 | 12.89 | 17.95 |
| *R<sub>merge</sub>* | 0.09638<br>(1.406) | 0.09237 (2.29) | 0.07939<br>(2.461) | 0.0782 (2.1) | 0.04741<br>(1.076) |
| *R<sub>meas</sub>* | 0.1027 (1.503) | 0.09991<br>(2.489) | 0.0859 (2.694) | 0.08437<br>(2.289) | 0.06705<br>(1.522) |
| *R<sub>pim</sub>* | 0.03461<br>(0.5238) | 0.03705<br>(0.9605) | 0.03182<br>(1.071) | 0.03117<br>(0.8963) | 0.04741<br>(1.076) |
| *CC<sub>1/2</sub>* | 0.999 (0.519) | 0.999 (0.305) | 1 (0.339) | 0.999 (0.333) | 0.999 (0.383) |
| *CC\** | 1 (0.827) | 1 (0.683) | 1 (0.711) | 1 (0.707) | 1 (0.744) |
| *Reflections used in refinement* | 86766 (8318) | 68444 (5443) | 73822 (7072) | 61857 (5563) | 36237 (3000) |
| *Reflections used for R-free* | 2000 (191) | 2000 (159) | 1998 (191) | 1999 (180) | 1999 (165) |
| *R<sub>work</sub>* | 0.1194<br>(0.2494) | 0.1310<br>(0.2538) | 0.1709<br>(0.3742) | 0.1306<br>(0.3122) | 0.1721<br>(0.3369) |
| $R_{free}$ | 0.1515<br>(0.2877) | 0.1696<br>(0.3032) | 0.2076<br>(0.4366) | 0.1651<br>(0.3540) | 0.2226<br>(0.4128) |
| $CC_{work}$ | 0.986 (0.768) | 0.983 (0.781) | 0.979 (0.615) | 0.982 (0.671) | 0.977 (0.675) |
| $CC_{free}$ | 0.985 (0.752) | 0.973 (0.722) | 0.973 (0.515) | 0.977 (0.653) | 0.973 (0.563) |
| Total number (N) of<br>non-hydrogen atoms | 8190 | 7932 | 7678 | 6441 | 5326 |
| N, macromolecules | 7919 | 7725 | 7513 | 6250 | 5204 |
| N, ligands | 21 | 26 | 21 | 21 | 23 |
| N, solvent | 250 | 181 | 144 | 170 | 99 |
| Protein residues | 279 | 279 | 285 | 285 | 285 |
| RMS (bonds) ($\text{\AA}$) | 0.007 | 0.023 | 0.004 | 0.004 | 0.002 |
| RMS (angles) ($^{\circ}$) | 0.98 | 1.74 | 0.75 | 0.70 | 0.51 |
| Ramachandran favored (%) | 96.57 | 96.55 | 95.33 | 96.92 | 97.26 |
| Ramachandran allowed (%) | 3.43 | 3.45 | 4.67 | 3.08 | 2.05 |
| Ramachandran outliers (%) | 0.00 | 0.00 | 0.00 | 0.00 | 0.68 |
| Rotamer outliers (%) | 1.42 | 0.48 | 1.63 | 0.60 | 0.73 |
| Clashscore | 2.90 | 3.24 | 5.23 | 2.61 | 1.37 |
| Average B-factor ($\text{\AA}^2$) | 12.69 | 14.33 | 13.82 | 15.14 | 21.52 |
| Average B-factor;<br>macromolecules ($\text{\AA}^2$) | 11.75 | 13.65 | 13.29 | 14.29 | 20.90 |
| Average B-factor; ligands<br>($\text{\AA}^2$) | 65.06 | 65.43 | 70.73 | 82.62 | 82.65 |
| Average B-factor; solvent ($\text{\AA}^2$) | 38.19 | 35.72 | 33.16 | 38.05 | 40.20 |

### 2.4 Data processing

Diffraction data recorded on Eiger 16M PAD detector (Casanas et al., 2016) was processed with the *XDS* package (Kabsch, 2010) and the programs *Pointless* (Evans, 2006) and *Aimless* (Evans and Murshudov, 2013), as implemented in the *autoxds* in-house processing script at SSRL (https://smb.slac.stanford.edu/facilities/software/xds/). Absorbed doses were calculated using *RADDOSE-3D (Bury et al., 2018; Zeldin et al., 2013)*.

## 3 Single conformer model refinement

Fig. 1 summarizes all refinement steps from the processed reflection data obtained above to the final multiconformer model. The first part of this process involves obtaining single conformer models via standard molecular replacement methods and iterative improvement of the model, which we briefly describe here.

**Fig. 1.**
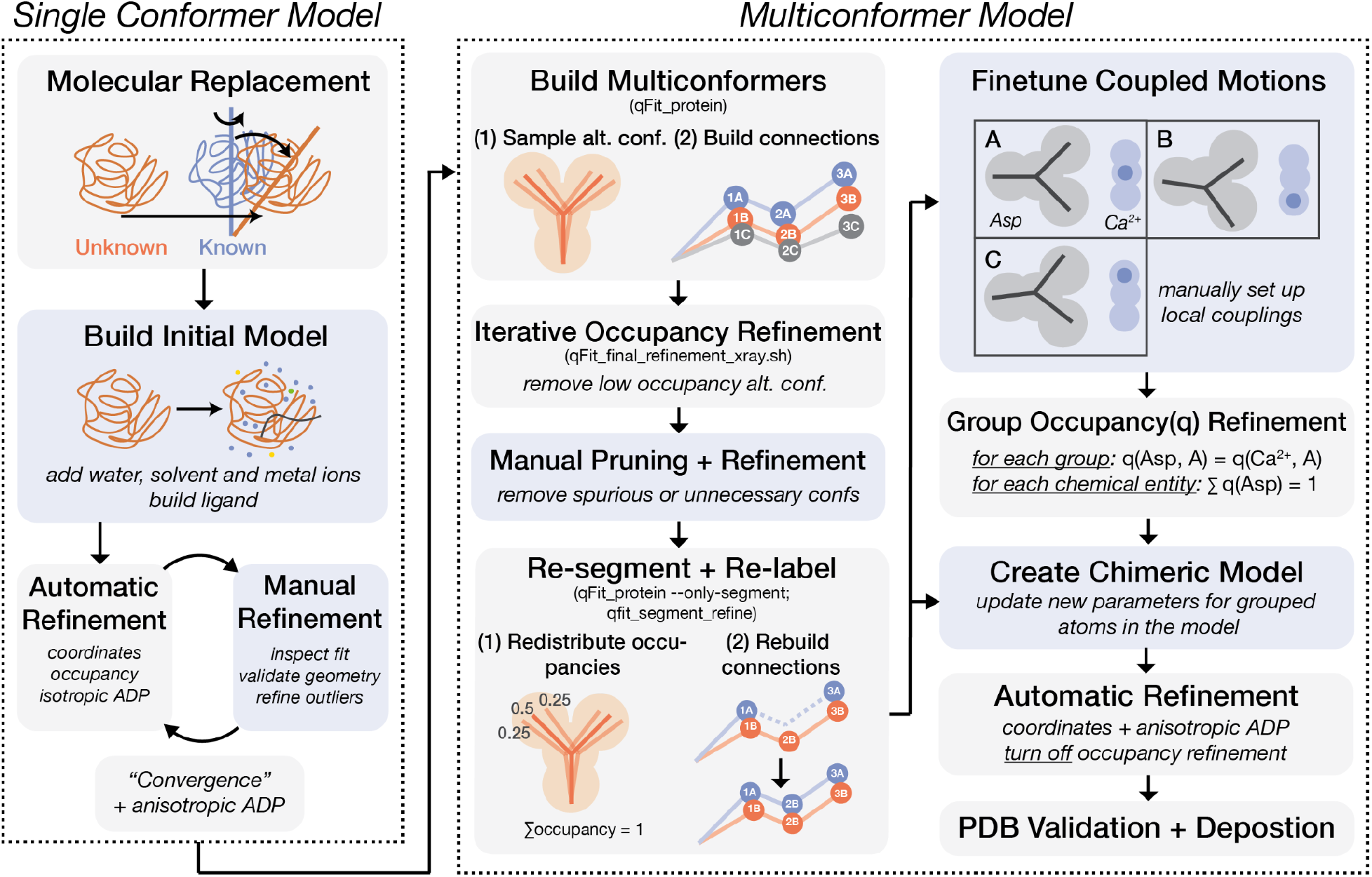
Flowchart for the refinement of a multiconformer model from diffraction data. Gray boxes indicate steps that are automated by refinement softwares such as *refmac* or *phenix*, or by *qFit* features; Blue boxes indicate steps that need manual interventions (e.g. in *Coot*). Abbreviations: ADP (atomic displacement parameter), alt. conf. (alternative conformations), q (occupancy).

### 3.1 Molecular replacement

Multiconformer modeling requires high quality data that is free of pathologies. These can be assessed using tools such as *phenix.xtriage* that can reveal the presence of twinning and translational noncrystallographic symmetry (tNCS). There are no pathologies in these high resolution Proteinase K datasets. Noting this, we proceeded to molecular replacement to obtain the initial phases (PDB: 3q5g; 100% sequence identity to wild type Proteinase K from *Parengyodontium album*). This search model was chosen because its crystallization was done in the same solvent as in our experiment. Molecular replacement (MR) was performed using the program *Phaser* after adding R_free_ labels to the reflection data.

### 3.2 Initial model building

We used the program *Coot* to examine the MR-generated model (.pdb) along with the density maps (.mtz), and manually complete an initial model. First, the C terminal carboxylate group was added to the model (using the *Add OXT at C terminus* tool) and the N- and C-terminus were refined (using *Real Space Refine Zone* and *Regularize Zone*). Next, we checked for the presence of any cis peptide bonds, as they are highly unfavorable (unless involving a proline residue) and may indicate model errors. One proline cis peptide bond was found for Proteinase K (as was present in the molecular replacement model) and was determined to be real as the model agrees with the 2F_o_ − F_c_ density map. Prior to refinement and after MR, we deleted all alternative conformers to obtain a single conformer model that is needed for later steps. We then cleaned up inorganic molecules (SO_4_, Ca^2+^) from the search model that are not present in our datasets. These molecules with no measured electron densities present were deleted, and other inorganic molecules were refined and edited such that each has occupancy = 1. Next, we added water molecules with electron densities above 1.4 rmsd using the *Find Waters* tool.

Before the refinement cycles, several simple validation metrics available in *Coot* were examined, including (1) Ramachandran plot, (2) geometry analysis and (3) rotamer analysis. Any outliers where atoms do not fit the densities well were refined using *Real Space Refine Zone* and *Regularize Zone*. Water molecules were examined using *Check/Delete Waters* where problematic water models were identified. In many cases, water molecules were too close to each other (< 2.4 Å), suggesting partial occupancies. These water pairs were edited so that they are alternative conformations of the same water molecule, and their occupancies were adjusted so that their combined occupancies do not exceed 1. Lastly, inorganic and water molecules were renumbered such that residue numbers are continuous within each chain.

### 3.3 Iterative model refinement

To improve the model and phases calculated from the model, it is necessary to perform multiple rounds of automatic refinement followed by manual adjustments until the convergence of a final single conformer model. In each initial round of automatic refinement (using the programs *refmac* and *phenix.refine*), 5 to 30 cycles of maximum likelihood refinement were performed for atomic coordinates, isotropic B-factors^1^ and occupancies; for final rounds of initial refinement, given the high resolution of the data, anisotropic B-factors were refined instead of isotropic. After each round of automatic refinement was completed, we manually inspected the F_o_ − F_c_ and 2F_o_ − F_c_ maps and the model in *Coot*. Difference (F_o_ − F_c_) map peaks above 5*σ* were examined in addition to the validation metrics mentioned above; any regions where the model did not match the 2F_o_ − F_c_ map were adjusted. Some of these peaks appeared to result from unmodeled alternative conformations and were expected to resolve after multiconformer modeling.

For the Proteinase K datasets, 5-6 iterations were performed until “convergence”. Here, we note that “convergence” is assessed remembering the adage that “refinement is never finished, but can be abandoned”. While one can continue the iterative refinement cycles infinitely, further improvements of model quality and agreement with experimental data will become lower in magnitude. Practically, we need to navigate these diminishing returns to determine whether we have arrived at a “final” model. We considered three aspects: (1) whether the models gave reasonable chemical representations of molecules, judged by the presence of outliers in torsion angles and geometry; (2) qualitatively, whether the model explains the density map well, judged mainly by the presence of interpretable F_o_ − F_c_ map peaks (above 4∼5*σ*) and how well the 2F_o_ − F_c_ map contours around the model; and (3) quantitatively, whether the measured structure-factor amplitudes |F_obs_| match the calculated amplitudes |F_calc_| from the current model, judged by R_work_ and R_free_ values (Brünger, 1992; Rupp, 2009). In these final single conformer models, a few outliers in backbone and sidechain torsion angles persisted, but they are likely real protein features as the model matches the 2F_o_ − F_c_ map shape. For example, D39, a member of the catalytic triad of Proteinase K, appeared to have unfavorable backbone torsions, and this outlier is not only observed in our datasets, but also in previously published PDB models. These regions where intrinsic conformational preferences are potentially perturbed by surrounding forces may be of interest for further investigation when modeling is complete – as they may arise from structural constraints or represent features that are evolutionarily-selected and provide functional benefits. All final single conformer models have R_work_ < 0.2, indicating a reasonably high model quality (Fig. 4B). R values appear to be larger for higher temperature datasets, which is expected due to increased thermal motions that cannot be accounted for by single conformer models.

### 3.4 Modeling an unknown ligand appearing at high temperatures

Intriguingly, at the Proteinase K active site, some unexplained electron densities gradually appeared for datasets obtained at higher temperatures. Because Proteinase K binds peptide substrates and the shape of these densities resemble a peptide chain, we reasoned that a short peptide may be able to bind at higher temperatures, and the apparent increase in the peptide density may reflect a shifted equilibrium favoring the bound state (Fig. 2A). While all the datasets were derived from crystals with the same content, the compositions of bound and unbound species in ordered parts of the crystals were different and generated different diffraction data and density maps, reflecting altered compositional heterogeneity across temperature. This heterogeneity information can be modeled by refining occupancies, as we described below.

**Fig. 2.**
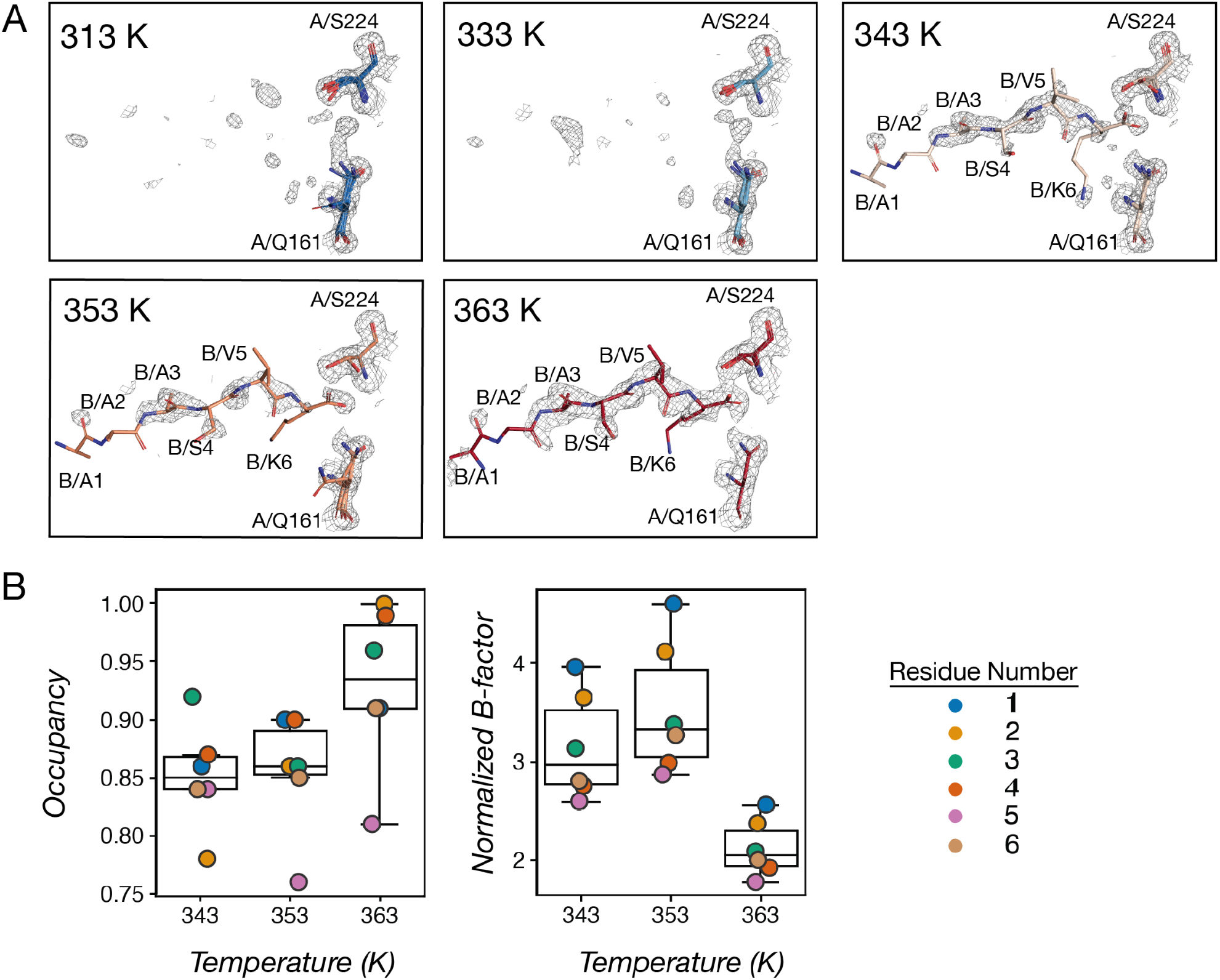
An unknown peptide is bound at the Proteinase K active site at high temperatures. **(A)** 2F_o_-F_c_ map and modeled residues for the binding site for datasets showed increasing electron densities for the bound peptide. Ser224 from the Proteinase K (chain A) is the catalytic serine, its backbone amide as well as Asn161 sidechain are the “oxyanion hole” hydrogen bond donors that interact with the carbonyl of the peptide ligand. The unknown peptides (chain B) were modeled for the 343, 353 and 363K datasets. **(B)** Occupancies and normalized B-factors for C*α* atoms of the unknown peptide residues.

To determine the sequence and the conformation of the unknown peptide, we used the 363K dataset which contains the most complete densities for this peptide as a guide. A poly-alanine chain (chain B) was built based on the overall 2F_o_-F_c_ density shape (using the *Add Terminal Residue* tool in *Coot*), followed by an automatic refinement round (using *phenix.refine*). Next, sidechain identities were estimated based on the shape of the 2F_o_-F_c_ densities that were not explained by the poly-alanine model; the F_o_-F_c_ map further inform sidechain choices (e.g. an unmodeled valine sidechain would give a signature shape of two adjacent negative density blobs). In *Coot*, non-alanine residues were mutated (using *Mutate & Auto Fit*), with the final sequence determined to be AAASVK. In the 343 and 353 K datasets, we modeled the same peptide sequence with roughly the same conformation as modeled in the 363 K dataset while fitting to local densities which are less complete than those in the 363 K dataset. Since the densities observed for this peptide are not complete, we set occupancies of chain B residues for all three datasets to a number below 1, which allowed the following automatic refinement step (using *phenix.refine*) to refine their partial occupancies (Fig. 2B). Overall, the occupancies of the peptide residues continue to increase from 343 to 363K, indicating higher bound species at higher temperatures. To compare the variability of these modeled positions across datasets, we calculated normalized B-factors by dividing the B-factors by the average B-factor of all atoms in each dataset. As expected, the peptide residues have higher-than-average B-factors due to incomplete densities (Fig. 2B). The normalized B-factors are lower for the 363K dataset, consistent with higher ordering of the bound species (Fig. 2B). In practice, there is some degeneracy between occupancy and B-factor refinement, but the refined results here, obtained from high resolution data, are consistent with greater occupancy and higher order (decreased B-factors) as temperature increases.

## 4 Multiconformer model refinement

Conformational heterogeneity from diffraction data can be represented by different metrics and data formats, each with its own limitations. In single conformer models, B-factors are typically used as a proxy for the degree of flexibility of a group, but they cannot be directly related to interpretable molecular geometries (e.g. atomic distances, bond angles and rotameric states) and involve contribution from other factors (e.g. crystallographic disorder) (Kuzmanic et al., 2014; Sun et al., 2019). Ensemble models generated using X-ray restrained molecular dynamics (MD) simulations provide 10s to 100s of separate single conformer models, where the relative population of different conformers reflect their occupancies (Burnley et al., 2012; Burnley and Gros, 2013; Forneris et al., 2014; Pearce and Gros, 2021; Ploscariu et al., 2021).

However, because of the high parameter-to-observation ratio, discrete conformers modeled for areas with ambiguous electron densities can be a result of overfitting instead of real conformational heterogeneity (Burling and Brünger, 1994; Wankowicz SA, 2020). Recent attempts to represent heterogeneity also include the use of bond-based parameters (bond lengths, angles and torsion angles) instead of cartesian coordinates; in this scheme, B-factors can be replaced by parameters describing the variation in torsion angles, which capture the physical nature of molecular motions more parsimoniously (Ginn, 2021). While promising in reducing the number of model parameters (and therefore reducing overfitting) and in improving the physical interpretability of X-ray models, refinement method based on this scheme (*Vagabond*) is still under development and has not achieved the accuracies of traditional Cartesian-based models by conventional R_free_ metrics (Ginn, 2021). In addition, both MD-based ensemble models and bond-based models are incompatible with current softwares for further manual or automatic refinements and therefore do not allow the fine-tuning of regions and structural features of interest, especially those detailing compositional heterogeneity that require more sophisticated refinement methods, which we described below (Section 4.4).

To improve interpretability, accuracy and compatibility while minimizing model complexity, we chose to refine each Proteinase K dataset into a multiconformer model, using the program *qFit* to initially sample and select alternative conformations (Keedy et al., 2015a; Riley et al., 2021; van den Bedem et al., 2009; van Zundert et al., 2018). In these multiconformer models, each protein residue has one to five alternative conformers, as needed to explain local densities; each conformer for a residue is assigned an “*altloc*” label (A, B, etc), and each atom for that conformer has its coordinates, occupancies, and B-factors recorded in a separate line in the model file. Unlike MD-based ensemble refinement, the approach we took only introduces additional parameters as needed to explain the experimental data; therefore, these models would be less likely to overfit. Practically, multiconformer models describe ensemble information in a single model following the conventional PDB (or mmCIF) format; thus, they are compatible with all common structural biology tools for further structural refinement and manual adjustment (e.g. in *Coot*) (Fig. 1). Both multiconformer models and MD-based ensemble models present visualization challenges. For example, in *PyMol* or *Chimera*, a multiconformer model is viewed in a single “state”, and the alternative conformations are all visible. For visualization, coloring by *altloc* id is helpful in interpreting coupled motions while viewing all modeled conformations. In contrast, ensemble models contain multiple “states” with whole copies of the entire system.

Scrolling through the states is helpful for visualization as viewing all models contained in the ensemble is often visually overwhelming. Further improvements in macromolecular visualization software for analyzing these complex model types will help further enable their use.

### 4.1 Automatic refinement using *qFit*

*qFit* is a Python-based software developed to automatically model and refine alternative conformers for protein residues and ligand molecules (Keedy et al., 2015a; Riley et al., 2021; van den Bedem et al., 2009; van Zundert et al., 2018). Here, we used *qFit* 3.0 to obtain initial multiconformer models for the Proteinase K datasets. To sample residue conformations, *qFit* first performs backbone sampling based on the anisotropic B-factors of the C_*β*_ atom (or O atom for Gly) which define the directionality of its potential motions, moving the atom around the ellipsoid while adjusting adjacent atoms (within a 5-residue segment) such that the backbone linkages are closed (Van Den Bedem et al., 2005). To sample sidechain conformations, *qFit* starts from the backbone conformations identified in the previous step and samples either around the C_*α*_-C_*β*_-C_*γ*_ bond for planar, aromatic sidechains or around the *χ* angles for other sidechains. At each *χ* angle and again once the entire sidechain is built, *qFit* evaluates the quality of sampled conformations and removes unnecessary and low occupancy conformers, keeping 1-5 optimal conformers for each residue whose positions, occupancies and B-factors best fit local densities.

After the optimal fitting of each individual residue, *qFit* reconnects the entire structural model taking account of conformer interconnections. Neighboring backbones residues with alternative conformations are split into segments, with each segment delimited by a residue with single-conformer backbone atoms. For each segment, *qFit* brings all residues to the same number of alternative conformations to avoid any “floating” conformers caused by missing backbones, and consistently assigns backbone occupancies and *altloc* labels (*qFit-segment*). Next, *qFit* determines the coupling of alternative conformers within each segment using a simulated annealing algorithm, relabeling all alternative conformers so that the coupled conformers do not clash (*qFit-relabel*). Details of the *qFit* algorithm have been described previously (Keedy et al., 2015a; Riley et al., 2021; van den Bedem et al., 2009; van Zundert et al., 2018) and the open-source software is available at https://github.com/ExcitedStates/qfit-3.0.

To obtain a multiconformer model from *qFit*, we need a single conformer model (.pdb) of reasonably high quality and a composite omit map (.mtz) (Terwilliger et al., 2008). A composite omit map provides the advantage of reducing model bias. To build such a map, the asymmetric unit is segmented into contiguous regions, and for the iterative refinement of each map region, model atoms located within that region are given an occupancy of 0 and therefore do not bias structure factor calculations; the final “composite” map then combines all refined segments (Terwilliger et al., 2008). A composite omit map was obtained from the single conformer model and map refined in section 3 using *phenix.composite_omit_map* with the *omit-type=refine* flag.

To sample conformers, we used the *qfit_protein* function with *-rmsd 0.1* setting, which removes redundant conformers when they have an all-atom RMSD below 0.1 Å. This RMSD setting was determined by testing *qfit_protein* with the default setting (RMSD threshold = 0.01 Å) and increased thresholds of 0.1 Å and 0.2 Å. Qualitatively, the 0.1 Å threshold produced the best model with a balance between conformation fit and parsimony. Setting an appropriate RMSD threshold in this step reduces model parameters and helps minimize manual efforts to prune conformers in later steps.

*qfit_protein* produced a multiconformer model (*multiconformer_model2.pdb*) that was then refined using the *qfit_final_refine_xray.sh* script. To ensure a parsimonious model, this refinement protocol involves iterative refinement (using *phenix.refine* functionalities) of atomic positions, occupancies and B-factors and removal of low occupancy (< 0.09) conformers until no such conformers emerge. This step produces a refined multiconformer model and map (with suffixes *_qFit.pdb* and *_qFit.mtz*).

### 4.2 Manual pruning and refinement

Manual inspection and refinement of the model and map from 4.1 are required for two reasons: (1) *qFit* may produce spurious conformers fitted to densities from noise or the bulk solvent and (2) additional backbone conformations may need to be added, as the backbone sampling of *qFit* 3.0 depends on the anisotropy of C_*β*_, which encodes backrub (Davis et al., 2006), crankshaft (Fadel et al., 1995; Fenwick et al., 2014), and shear (Hallen et al., 2013; Smith and Kortemme, 2008) motions, but does not report on large backbone rearrangements such as the 180° peptide flips (Keedy et al., 2015a).

In *Coot*, we inspected each residue to prune any spurious or unnecessary conformers, including those that do not fit to local densities, those that would cause strain or clashes with neighboring residues, and those that are too similar. While the criteria for similarity may be qualitative and *ad hoc*, we note that both sidechain and backbone atoms need to be compared to decide if a conformer needs to be pruned. For example, two conformers may have the same sidechain conformation but obviously different backbone positions. In this case, both sidechain conformers need to be kept in the model, as the current PDB format will not allow two sets of backbone atom positions linked to only one sidechain conformer (even though a single backbone conformation can spawn two side chain conformations). The sidechains in solvent-exposed areas are more likely to show spurious conformers. In some cases, there were no 2F_o_-F_c_ contours even at < 0.5 *σ* around the spurious conformers and also no positive F_o_-F_c_ peaks, suggesting that these conformers may have been incorrectly fitted to densities resulting from noise or bulk solvent contributions. Only the conformers supported by the 2F_o_-F_c_ map were kept in the model. In the meantime, we checked for any backbone conformations that needed to be rebuilt or sidechains that could be refined to fit the 2F_o_-F_c_ map better, manually adjusting their positions as needed.

### 4.3 Automatic relabeling of structural segments

Manual pruning and refinement are essential to correct and improve the model, but also introduce model inconsistencies that need to be resolved. First, because some conformers were deleted, the combined occupancies of the remaining conformers of a residue did not sum to one. Second, deletion of conformers resulted in breaks in peptide linkages. To redistribute occupancies and reconnect the peptide, we re-ran *qfit_protein* with the flag *–only-segment.* With this option, *qFit* does not re-sample and score residue conformers, but re-distributes the occupancies of the remaining conformations and performs the segmentation and labeling step as described in 4.1 (*qFit-segment* and *qFit-relabel*). This step is followed by another automatic refinement cycle using *qFit_segment_refine*.

This procedure generates connected backbones with consistent occupancies for coupled neighboring conformers, but at the cost of increased number of parameters, as it requires bringing in duplicate conformers. For example, if residue N has four alternative backbone conformations (A, B, C, D) and residue N+1 has two alternative conformations (A, B), this procedure will create C and D conformers for residue N+1 by duplicating its A and B conformers. This duplication may continue until we reach the end of a segment, so that all backbones have the same number of alternative conformations (A, B, C, D) and are therefore properly connected. The alternative to the duplication of conformers is to have “floating” backbone atoms, e.g. with residue N conformers C and D having no connection from the backbone carbonyl to the next alpha carbon. Ideally, we would like to have a nested model format where the C and D conformations can be “children” of the A and B conformations, but currently, neither the PDB nor CIF format currently allow for that representation (Hancock et al., 2022; Pearce et al., 2017; Vallat et al., 2023).

### 4.4 Modeling coupled motions

To finetune the model for regions of interest where coupled motions may occur, we used a constrained group occupancy refinement approach, which we illustrate below using the example of the Proteinase K Ca^2+^ binding site. This binding site was identified in previous structural studies (Betzel et al., 1988), where a Ca^2+^ ion is coordinated by the sidechain of D200, the backbone O atoms of V177 and P175, and surrounding water molecules (Fig. 3A).

**Fig 3.**
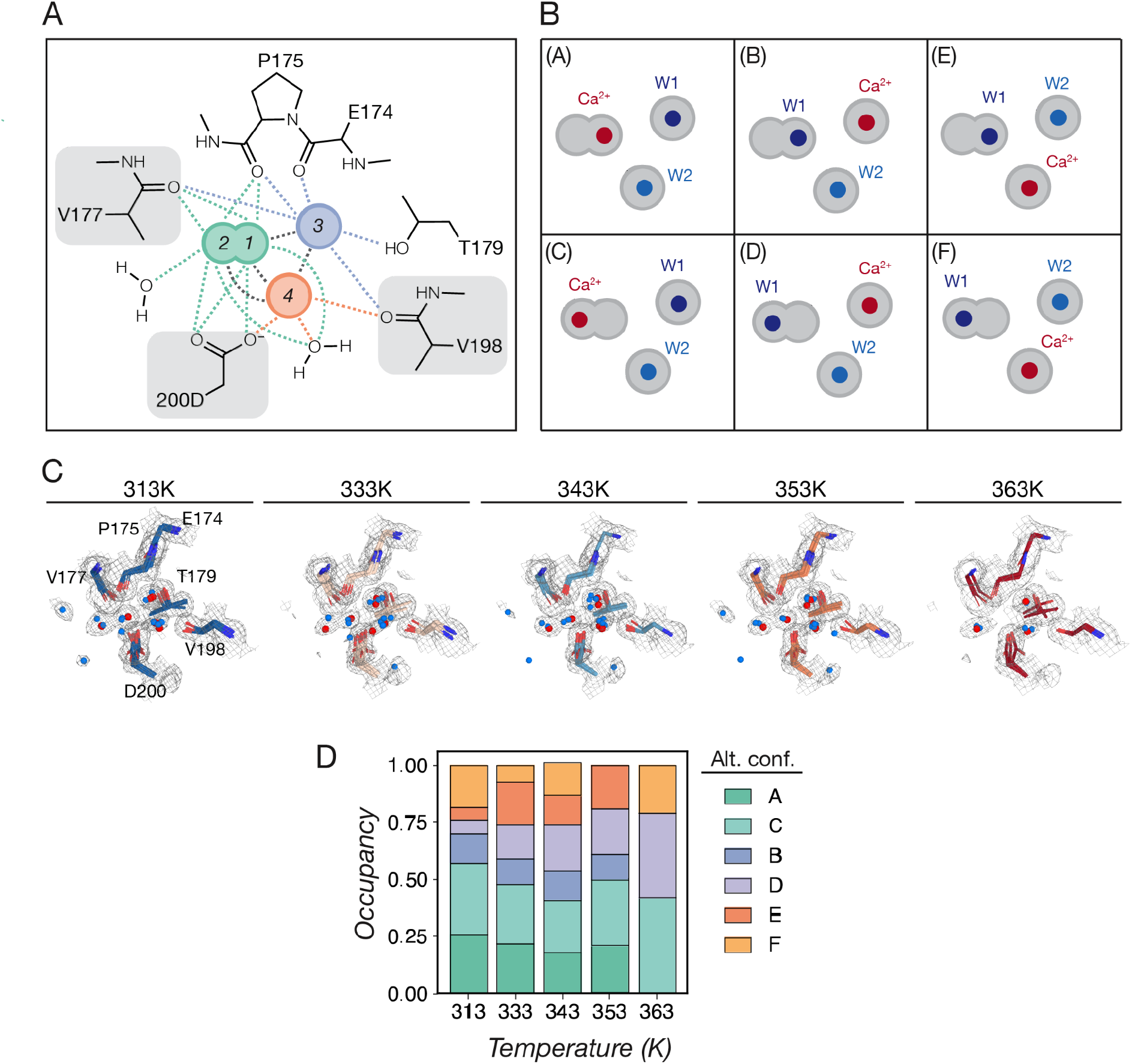
Modeling and refinement of the Proteinase K Ca^2+^ binding site. **(A)** Four possible positions for Ca^2+^ as suggested by the 2F_o_-F_c_ maps; each position appears to be stabilized by 5 to 7 metal-coordinating interactions with surrounding water molecules or protein residues. Position 1 corresponds to the Ca^2+^ position that is typically modeled. Protein residues that showed alternative conformers orienting towards different Ca^2+^ positions are indicated by gray boxes; these residues were included in the group occupancy refinement. **(B)** All possible configurations (conformers A through F) for the Ca^2+^ and water molecules. **(C)** 2F_o_-F_c_ map and models for the Ca^2+^ binding site, including all alternative conformations for each model. Water molecules are shown as blue spheres, and Ca^2+^ as red spheres. **(D)** Changes in the refined occupancies of each alternative conformation across temperature.

Nevertheless, our diffraction data revealed a more complex picture for Ca^2+^ interactions in this binding site: the 2F_o_-F_c_ map does not clearly indicate one unique position for the Ca^2+^; instead, for datasets obtained at 313 to 353K, there are four spherical densities within this binding site, and two of these spheres are very close together, with their merged densities forming a dumbbell shape (Fig. 3C). Our density map and initial multiconformer model suggested that the Ca^2+^ can occupy these alternative positions in the binding site, for three reasons. First, the commonly modeled position where Ca^2+^ forms a bivalent interaction with the D200 sidechain lies within the overlapping dumbbell-shaped density, suggesting alternative position instead of a coordinating water molecule, as the interaction distance would be too close (<2 Å) and highly unfavorable (Fig. 3C). Second, the alternative conformers modeled for nearby residues such as D200 and V177 include those that orient towards positions other than the commonly modeled one, suggesting that these residues can stabilize Ca^2+^ when it occupies these other positions (Fig. 3C). Lastly, in the 363K dataset, densities for the commonly modeled position disappeared and the dumbbell-shaped density shrinked to an elliptical shape, suggesting that alternative conformations are favored at high temperatures (Fig. 3C, D). To unambiguously determine possible positions of Ca^2+^, future experiments can collect diffraction data at longer wavelengths to detect Ca^2+^ anomalous signals; here, we considered all possible alternative configurations as suggested by the electron density maps.

To model alternative positions of Ca^2+^ and how the interacting residues move accordingly, we manually set up alternative conformers of Ca^2+^ and its surrounding protein residues and water molecules as “groups” in *Coot* by creating multiple copies of the same atoms and assigning the same *altloc* label to the atoms in the same configuration (Fig. 3B). Then, we used *phenix.refine* to perform automatic refinement with group occupancy constraints that will produce consistent occupancies for chemical entities within a group.

To enumerate all possible Ca^2+^ and binding site residue configurations, we considered each of the four spherical densities as potential alternative positions for Ca^2+^, and in each case assigning the other density blobs as water molecules. Because only one atom can occupy the dumb-bell region at a time, we modeled one Ca^2+^ and two water molecules for each alternative conformation. In total, there are 6 different configurations for the Ca^2+^ and the two coordinating waters as a group, as illustrated in Fig. 3C; we therefore created alternative conformations A through F for these molecules accordingly. Next, we identified protein residues that showed correlated motions with these different Ca^2+^ positions, which include D200, V177 and V198 (Fig. 3A, B). We also modeled 6 alternative conformers (A through F) for each of these residues, and their alternative conformer labels were reassigned so that each conformer was in the correct group. For example, the conformer of D200 that is the closest to the A conformer of Ca^2+^ was labeled “A”, et cetera. For the 363 K dataset, the dumbbell-shaped density observed for other datasets diminished into an eclipse with no clear indication for two separate configurations (Fig. 3C); therefore, duplicate configurations (A, B and E) were removed.

To model the positions, B-factors and occupancies of these chemical entities as a group, we included group occupancies refinement strategies in our next cycle of *phenix.refine*, assigning each alternative configuration as a constrained group (e.g. group A was the A conformers of Ca^2+^, waters, D200, V177 and V198). Using this approach, all atoms in a group are refined to the same occupancy and each chemical entity will have a total occupancy of 1 summed over all its alternative conformers. The positions and B-factors were also allowed to further refine.

Nevertheless, occupancy refinement would be performed for the entire model, and some alternative conformers of protein residues outside the Ca^2+^ binding site may drop below 0.09 again. Therefore, we created chimeric models that merged the pre-grouped model (from **4.3**) with the updated positions, B-factors and occupancies for grouped atoms in the grouped and refined model obtained here. Additional refinement runs for this chimeric model were then performed with fixed occupancies, allowing only the atomic positions and B-factors to fluctuate.

Increasing temperatures favor alternative Ca^2+^ binding configurations, as suggested by the refined group occupancies: at lower temperatures, Ca^2+^ mainly occupies the dumbbell region (configurations A and C); as temperature increases, the occupancy for Ca^2+^ at the more distal position (configurations B and D) increases (Fig. 3D). In addition, the elliptical instead of dumbbell shape for the center density at 363K suggests less distinction and therefore higher mobility for exchanging between the A and C sites (Fig. 3C).

Overall, the final multiconformer models showed decreased R factors across all datasets (Fig. 4), indicating improved fit of the models to the underlying data after multiconformer refinement; in particular, the decrease in the cross-validation term R_free_ suggests that the improved accuracy does not arise from overfitting (Fig. 4A).

**Fig 4.**
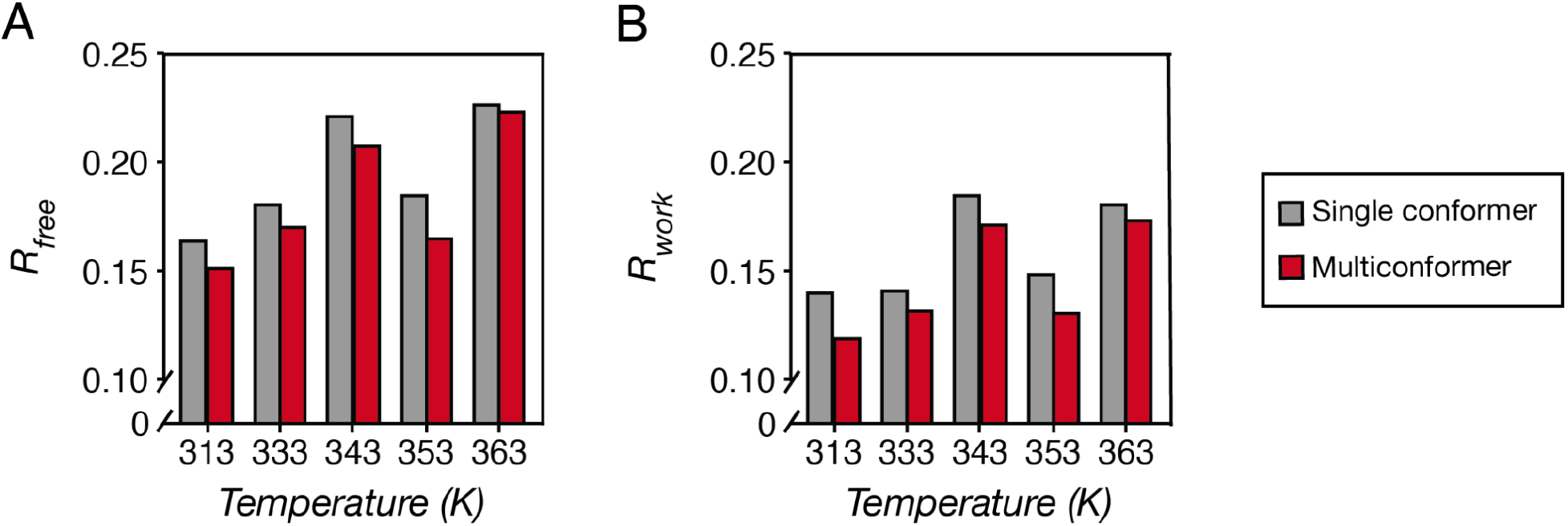
R_free_ **(A)** and R_work_ **(B)** for the final single conformer *v.* multiconformer models indicate improved model accuracy after multiconformer refinement.

## 5 Identifying temperature-dependent conformational changes

Multiconformer models provide rich information for protein conformational ensembles, but it can be difficult to extract conformational changes that are significant and relevant to functional aspects of interest. This difficulty arises from the fact that each residue may have a different number of alternative conformers modeled for different datasets, and each alternative conformer has their own modeled positions, occupancies and B-factors, preventing a matched statistical comparison across datasets. Here, we used the program *Ringer* (Lang et al., 2010) to guide our search for interesting conformational changes and identified widespread changes of the proteinase K ensemble in response to temperature. The approach that we describe here can also be extended to study other structural perturbations, such as ligand binding and mutations.

### 5.1 Ringer analysis

The *Ringer* program systematically samples electron densities around sidechain rotamers, allowing for the detection of low-occupancy sidechain conformational states and the comparison of sidechain states across datasets at different temperatures. *Ringer* analysis complements multiconformer models, as it provides torsional electron density profiles for all sidechains at 5 or 10° intervals that can be systematically compared across datasets. Nevertheless, *Ringer* can only sample electron densities around sidechains based on backbone positions from a single conformer model; thus, the resulting profile reflects a mixture of sidechain and backbone motions. For example, a broad *Ringer* peak may result from a highly flexible sidechain attached to constrained backbone atoms, or the opposite, or a moderate level of flexibility from both. Therefore, to distinguish between these possibilities, one must return to the multiconformer model and electron density maps.

*Ringer* can be accessed via *phenix* using the *mmtbx.ringer* command with a model and a single conformer model supplied (*mmtbx.ringer model.pdb map.mtz*). We used the single conformer model from **3.4** and the final map after multiconformer refinement from **4.4**, as the final map provides more accurate electron densities. This command produces a table of electron densities for each residue-rotamer in the model from 0 to 359° at specified intervals (default 5°).

The raw *Ringer* profiles are helpful for the interpretation of weak densities and further refinement of the multiconformer model. Crudely, any rotamer angles at ≥ 0.3 *σ* are likely to be conformational features rather than noise from hydrogens (Lang et al., 2010). One may return to the multiconformer model to refine particular areas as informed by *Ringer*.

For the systematic comparison of rotamers dynamics across datasets, we need to normalize *σ* values (eq. 1), as the scale of electron density values can vary across datasets and obscure changes of *σ*.

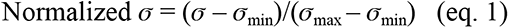

The normalized *Ringer* profiles revealed diverse patterns of sidechain conformational changes across temperatures. In the simplest scenario, we would expect high temperatures to favor higher-entropy states, and the distributions of sidechain rotamer angles are expected to broaden. One example of this pattern is the *χ*1 of Glu43, as indicated by the broader shoulders of the 363K *Ringer* profile and its more dispersed 2F_o_-F_c_ densities around the sidechain (Fig. 5A). In the second case, we observed the emergence of an alternative sidechain rotamer at higher temperatures, such as for the *χ*1 of the catalytic residue Ser224 (Fig. 5A). Unexpectedly, we also observed the disappearance of rotamer states at high temperatures, such as for the *χ*1 of Ser63, emphasizing the idiosyncrasy of temperature effects on individual rotamers, residues and regions, rather than a universally higher flexibility (Fig. 5A). Lastly, there are also highly positioned residues such as Asn163 whose *χ*1 profiles do not change across temperature (Fig. 5A).

**Fig. 5.**
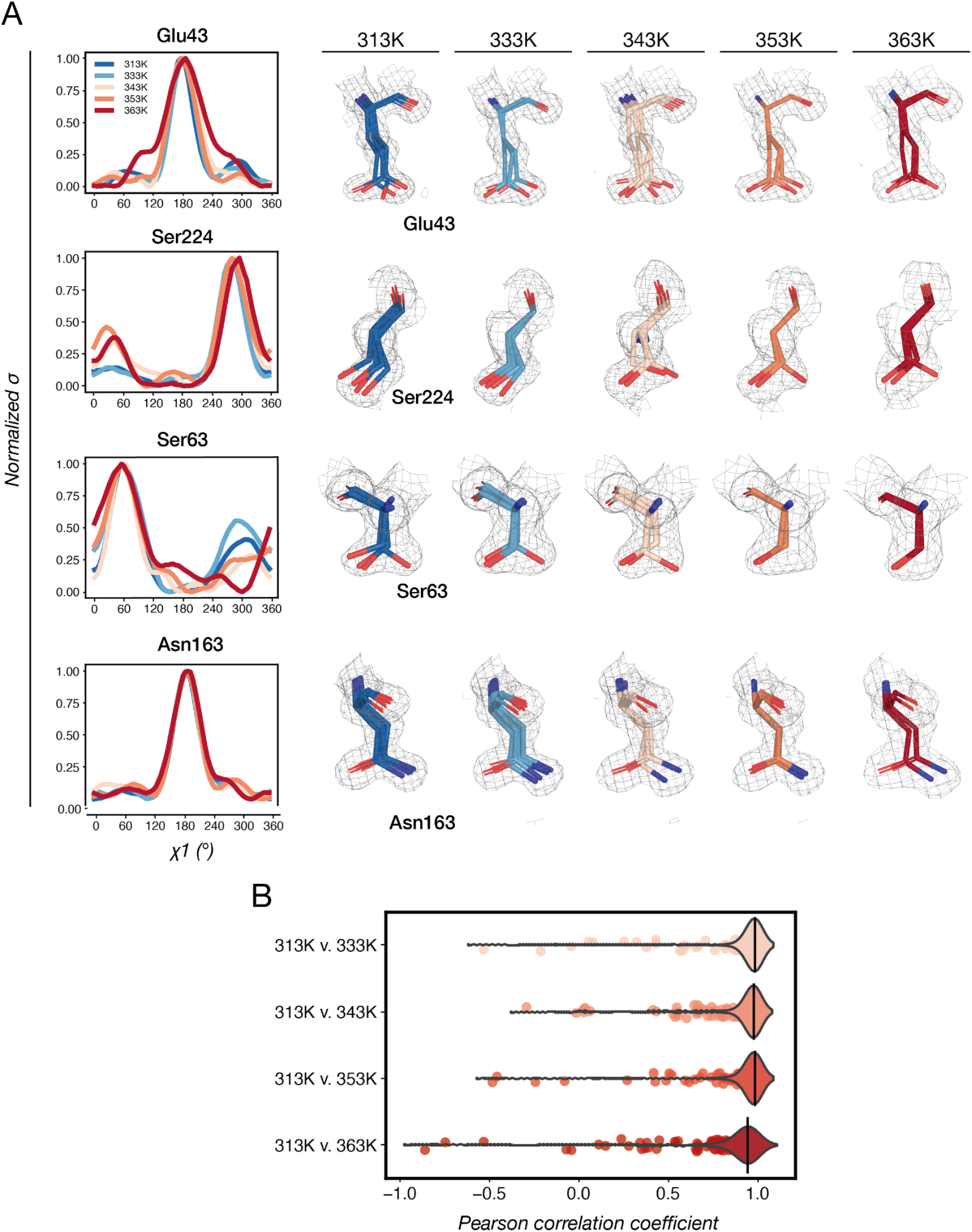
Temperature-dependent rotamer changes in Proteinase K. **(A)** Examples for how rotamers change across temperatures and their *Ringer* profiles. 2F_o_-F_c_ maps are contoured at 1*σ*. **(B)** Distributions of P_CC_ for comparisons of all rotamers across datasets.

To further quantify the similarities and differences between *Ringer* profiles, we calculated Pearson correlation coefficients (P_CC_, or Pearson’s *r*) using the *scipy.stats.pearsonr* function of the *SciPy* package (Virtanen et al., 2020). Across sidechain rotamers of the entire Proteinase K structure, P_CC_ values decrease when comparing the 313K model to higher temperature models, and are especially low for the 363K dataset (Fig. 5B). For the comparison of 313K *versus* 363K dataset, we identified 86 rotamers with P_CC_ ≤ 0.9 among a total of 410 rotamers, suggesting widespread conformational changes in response to higher temperatures. As all datasets here were collected above the glass transition, these changes are mostly subtle, and we would expect more significant changes for comparisons of datasets below and above glass transition (Fraser et al., 2011; Halle, 2004; Keedy et al., 2014; Rasmussen et al., 1992; Tilton et al., 1992).

## 6 Summary and Conclusions

Conformational ensembles, rather than static structures, are needed to deepen our understanding of protein functions and ultimately reach the ability to derive quantitative, predictive models for protein functions (Austin et al., 1975; Benkovic et al., 2008; Benkovic and Hammes-Schiffer, 2003; Frauenfelder et al., 1991, 1988; Hammes et al., 2011; Mokhtari et al., 2021) . Nevertheless, X-ray derived ensemble data is limited due to experimental challenges [which we addressed in (Doukov et al., 2020)] and the requirement for specialized refinement approaches, which are not easily accessible. Here, we used a series of Proteinase K datasets collected at increasing temperatures to provide a practical and detailed tutorial for the refinement of multi-conformer models and correlated motions within these models, and we discussed the rationale behind our refinement choices and their advantages and limitations. We note that many of our refinement choices are limited by the PDB format and interpretations by refinement softwares. In particular, multiconformer models need to account for alternative conformations for each individual residue as well as the connections between conformers across the protein backbone, and this multidimensional information cannot be cleanly represented by the “flat” PDB format without duplicated model parameters. The mmCIF format could potentially represent multiconformer connectivities and interrelationships because of its more flexible formatting. Such future efforts will need to evolve with projects that have high compositional [e.g. fragment screening (Krojer et al., 2020; Weiss et al., 2022)] and conformational [e.g. time-resolved serial femtosecond crystallography (Oda et al., 2021; Schmidt, 2021)] heterogeneity. Meeting these challenges will also help build molecular models compatible with increasingly complex 3D classification and heterogeneous map reconstruction methods in cryo-EM (Zhong et al., 2021). In addition, we encountered issues during refinement and PDB deposition because many widely-used tools (e.g. *MolProbity* and *Reduce*) are not optimized for multiconformer models. We suggest that future efforts in improving the PDB/mmCIF format and structural biology tools to accommodate ensemble features will simplify the process of obtaining ensemble models and allow the database of conformational ensembles to grow.

Proteinase K appears to undergo widespread conformational changes across temperature. These observed changes are potentially linked to its stability, binding and catalysis, such as the increased occupancies of the bound peptide ligand, changes in Ca^2+^ binding configurations, and altered distributions of rotameric angles for catalytic residues. While *qFit* automates the sampling of alternative conformations and provides a preliminary model, we emphasize that additional finetuning is needed to improve the accuracy of the model and to extract interesting local changes. For example, we showed that the Ca^2+^ binding site can be modeled by 6 different alternative configurations and determined how the occupancies of each change across temperatures. This strategy may be extended to model other coupled motions of interest, e.g. to determine if the motions of the active site groups are constrained or facilitated by surrounding residues, or if the binding of an allosteric ligand shifts the equilibrium of conformational states of a network of residues that move together. We expect that these modeled changes will lead to hypotheses that can be tested by additional experiments – for example, by introducing structural perturbations that change the magnitude or direction of these motions or disrupt their couplings.

## Acknowledgements

This work was funded by a National Institutes of Health (NIH) grant (GM145238) to J. S. F. and a National Science Foundation (NSF) grant (MCB-1714723) to D.H. Use of the Stanford Synchrotron Radiation Lightsource (SSRL), SLAC National Accelerator Laboratory is supported by the US Department of Energy, Office of Science and Office of Basic Energy Sciences under Contract No. DE-AC02- 76SF00515. The SSRL Structural Molecular Biology Program is supported by the DOE Office of Biological and Environ- mental Research and by the National Institutes of Health (NIH), National Institute of General Medical Sciences (NIGMS, P41GM103393). The contents of this publication are solely the responsibility of the authors and do not necessarily represent the official views of NIH or NIGMS. We thank Lisa Dunn (SSRL) for help with scheduling experimental beam time.

## Footnotes

1 B-factors are also named thermal factors, temperature factors or atomic displacement parameters (ADP) and used interchangeably.

